# Metabolic fluctuations explain allometric scaling diversity

**DOI:** 10.1101/2025.11.06.686202

**Authors:** Andrea Tabi, Wout Merbis, Fernando A Nobrega Santos, Ricard Solé

## Abstract

Metabolic scaling, the relationship between energy use and body size, has long been treated as a universal law of life. However, extensive variation in scaling exponents across species challenges this assumption. Here, we show that such scaling can emerge spontaneously from stochastic cellular growth dynamics, without postulating any fixed relationship between mass and metabolism. In our framework, ontogeny is a nonequilibrium thermodynamic process in which energy is continuously dissipated and redistributed among fluctuating cellular states. When applied across diverse life histories, the model reproduces the observed range of metabolic exponents, revealing that scaling diversity arises naturally from fundamental thermodynamic constraints on stochastic, energy-dissipating growth.

## I INTRODUCTION

The quest for universal patterns in biology has long attracted scholars from various disciplines, including physics, mathematics, and evolutionary biology [7, 15, 19, 31, 45]. Several general principles have emerged from the study of fractals and geometric scaling laws [30], network topology and dynamics [5], criticality and fluctuations [8, 14, 33, 41, 43], emergent phenomena and evolutionary constraints [2, 39, 40]. Among these unifying frameworks, perhaps the most influential in the biological sciences is *metabolic scaling theory* (MST) [6, 44]. MST has shaped ecological theory for several decades in its effort to unify processes from individual physiology to ecosystem dynamics. Its central premise is that an organism’s metabolic rate scales predictably with its body mass and environmental temperature. Specifically, metabolic rates are assumed to follow the Boltzmann-Arrhenius kinetics with respect to temperature and scale sublinearly with mass according to *Kleiber’s law* [23]:

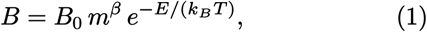

where *B* is the metabolic rate, *m* is body mass, *T* is absolute temperature, *B*_0_ is a normalization constant, *β* is the *allometric exponent* (typically taken as 3*/*4), and *E* is an *activation energy*, generally ranging between 0.2 and 1.2 eV.

Empirical studies have revealed substantial variation in allometric exponents both within and across taxonomic groups. Reported values range from highly superlinear (*β* ∼ 1.7–2.0) in prokaryotes, to nearly linear (*β* ∼ 1.0–1.1) in protists, and sublinear (*β* ∼ 0.76–0.79) in metazoans [10, 11, 21]. These discrepancies raise fundamental questions about the existence of a truly universal metabolic scaling law.

A wide variety of theoretical models have been proposed to explain the diversity of observed metabolic scaling relationships [1, 17]. These models are based on various mechanisms, such as chemical engineering principles, to explain the variety among small aquatic animals [36], using the surface-to-volume ratio of the organism assuming a constant body density as the true value [20, 37], elastic similarity based on the energy turnover of muscle contraction [32], combination of gravitational force and surface area [13] or cell size [25].

One of the most influential explanations is based on the geometry of distribution networks, such as vertebrate cardiovascular and respiratory systems, plant vascular systems, and insect tracheal networks, which proposes that the 3*/*4 power law arises from their hierarchical, fractal-like organization filling space [4, 46]. While widely accepted, this framework has been extensively debated, with critics highlighting both empirical inconsistencies and theoretical oversimplifications [12]. Most recently, ontogenetic and energy budget models have sought to recover the 3*/*4 exponent by incorporating life-history optimization principles [22, 29, 49]. In a simple thermodynamic framework, the variation in metabolic scaling exponents is explained as a balance between the energy dissipated as heat and the energy efficiently used, however the framework still relies on assumed power-law relations rather than deriving them from first principles [3].

Here we go beyond this by deriving scaling exponents and its variability from stochastic cellular-level fluctuations, without imposing scaling laws *a priori*. We present a stochastic ontogenetic model, where we apply thermodynamic principles to show that allometric scaling emerges from cellular level fluctuations and their thermodynamic costs. Our approach sheds light on individual and species-level variations in allometric scaling exponents as well as provides a mechanistic explanation as to how allometric scaling emerges.

## II. METHODS

### A. Stochastic ontogenetic growth model

Our ontogenetic growth model integrates principles of energy conservation at the cellular scale with stochastic fluctuations. Unlike classical ontogenetic models, our framework captures individual-level variability by incorporating stochastic cell dynamics, providing a more realistic representation of individual growth development. Individual growth is modeled as an emergent outcome of energy allocation among maintenance, biosynthesis, and heat loss that account for the stochastic fluctuations in cell dynamics. We assume that individuals have a genetically predetermined maximum number of cells (*N*_max_), sampled from a log-normal distribution characterized by a species-specific mean 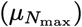 and variance 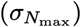. The trajectory of cellular growth follows a stochastic logistic equation with a stochastic Laplace distributed term that is based on empirically observed non-Gaussian Laplace-like noise in the metabolic rates of the organism [27]:

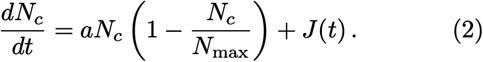

Here *J* (*t*) ∼ Laplace(0, *b*) is the stochastic term, mimicking bursts of proliferative activity or cellular perturbations with state-dependent scale parameter (*b*). This is proportional to the number of cells and decreasing with developmental progress (see SM for details):

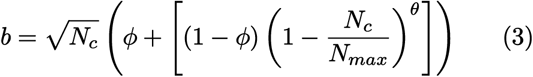

where *ϕ* is the baseline fluctuation level and *θ* sets the nonlinearity (in simulations we set it *θ* = 2). We assume that fluctuations arise independently in each cell, i.e. the total fluctuation magnitude grows as 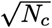. The rate of fluctuations decays over time, reflecting decreasing metabolic variability in maturity. The inclusion of such a term allows for a more realistic accounting of metabolic energy usage during development, especially in tissues undergoing rapid or irregular cell turnover. Mechanistically, the presented growth model can be derived from a microscopic birth-death process (see SM for details). The difference of these two independent processes results in a double exponential (Laplace) distribution. Traditional models of ontogenetic growth often assume that energy used for biomass accumulation is converted with near-perfect efficiency, neglecting the role of heat dissipation (Table I). However, mounting empirical evidence suggests that developmental processes are energetically costly and far from thermodynamically optimal [51]. Thus, the total energetic expenditure in our model has three major components based upon energy-expenditure budget theory [24]:

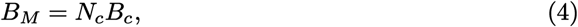

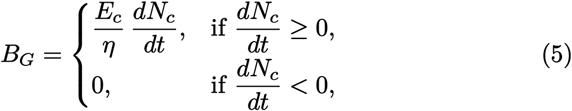

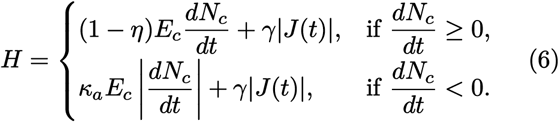

**Table 1.**
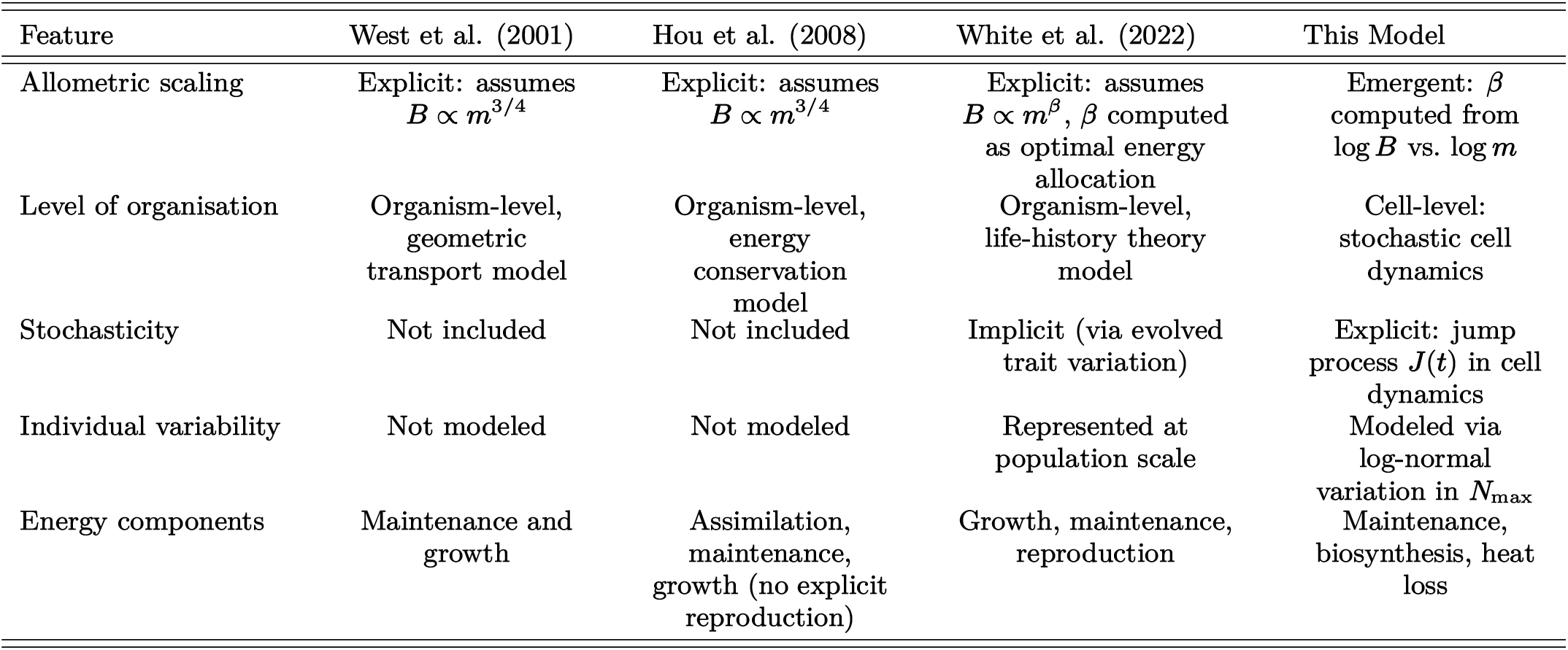
Comparison of ontogenetic growth models.

| Feature | West et al. (2001) | Hou et al. (2008) | White et al. (2022) | This Model |
| --- | --- | --- | --- | --- |
| Allometric scaling | Explicit: assumes $B \propto m^{3/4}$ | Explicit: assumes $B \propto m^{3/4}$ | Explicit: assumes $B \propto m^\beta$, $\beta$ computed as optimal energy allocation | Emergent: $\beta$ computed from $\log B$ vs. $\log m$ |
| Level of organisation | Organism-level, geometric transport model | Organism-level, energy conservation model | Organism-level, life-history theory model | Cell-level: stochastic cell dynamics |
| Stochasticity | Not included | Not included | Implicit (via evolved trait variation) | Explicit: jump process $J(t)$ in cell dynamics |
| Individual variability | Not modeled | Not modeled | Represented at population scale | Modeled via log-normal variation in $N_{\max}$ |
| Energy components | Maintenance and growth | Assimilation, maintenance, growth (no explicit reproduction) | Growth, maintenance, reproduction | Maintenance, biosynthesis, heat loss |

where *B*_*M*_ is the cellular maintenance term, *B*_*G*_ is the growth-energy term, and *H* is the metabolic inefficiency term. Here *B*_*c*_ is the per-cell maintenance cost, *E*_*c*_ the energy to build a cell, *η* ∈ (0, 1] the growth efficiency, *κ*_*a*_ the apoptosis cost (set to 0.3) and *γ* the coefficient converting stochastic activity into heat. The cost of maintaining cells depends on the number of cells (*N*_*c*_) and the basal metabolic rate per cell. The energetic costs of growth is calculated as the average energy required to create a new cell (*E*_*c*_) and the change in cell numbers (*dN*_*c*_*/dt*). The metabolic inefficiency (*H*) consists of the stochastic term modulated by the stochastic inefficiency coefficient (*γ*) and the inefficiencies during growth phase. This additional energetic cost captures inefficiency associated with stochastic bursts of cell proliferation or loss. This energy is not used for biosynthesis, but is dissipated as heat or transient metabolic overhead, reflecting the energetic consequences of cellular stochasticity.

The organismal mass evolves as [47]:

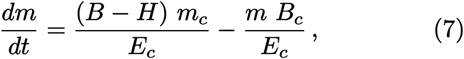

where *m*_*c*_ corresponds to the mass per cell. At steady state, we can derive analytical expressions for the expected adult metabolic rate, i.e.:

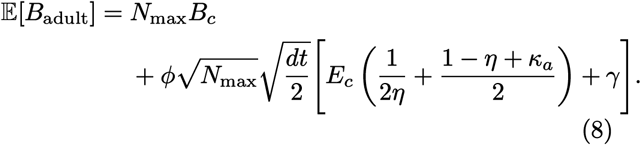

which combines the baseline cellular maintenance cost (*N*_max_*B*_*c*_) with additional energetic contributions from biosynthesis, metabolic inefficiency, and stochastic fluctuations. The expected adult mass is

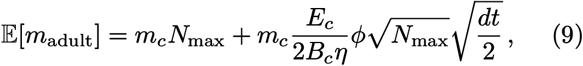

given by the sum of the total cellular mass at carrying capacity and an additional term arising from stochastic energy allocation during growth, which asymptotically converges to zero as the maximum number of cells increases. Using this, the effective allometric scaling exponent:

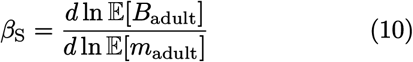

quantifies the scaling relationship between metabolic rate and body mass at steady state, providing a theoretical prediction in the presence of cellular stochasticity (see SM for details). These relations link the scaling of metabolic rate and adult mass to underlying cellular stochasticity, demonstrating how energy conservation principles and metabolic fluctuations jointly shape ontogenetic trajectories. The scaling exponent naturally emerges from the combination of deterministic and stochastic components of the growth model (see SM for details).

In our framework, ontogenetic growth is a nonequilibrium thermodynamic process involving cellular turnover and thermodynamic dissipation. Stochastic cellular turnover includes cellular activities such as proliferation, maintenance, and death. In organismal development, these processes are not perfectly efficient, but a portion of stochastic bursts of cell proliferation or loss inevitably dissipates as heat [51]. These fluctuations are expected to exhibit a non-Gaussian mixed distribution with exponential tails that show most similarities with Laplace distribution. We quantify the magnitude of metabolic fluctuations (*r*) in steady state (adult phase), namely

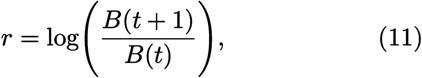

with the mean absolute deviation of the Laplace distribution (*σ*_*r*_):

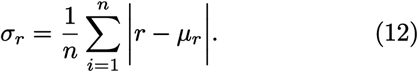

We have an analytical form to estimate *σ*_*r*_ only if jumps are very small (for details see SM). At larger jumps, however, the analytic expression overestimates the variance, thus we estimate *σ*_*r*_ using Monte Carlo sampling (see SM for more details). When the jump process dominates the metabolic fluctuations in adult phase, *σ*_*r*_ saturates at a universal limit as a consequence of logarithmic compression 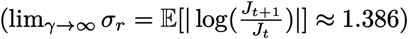.

A key distinction between our approach and previous ontogenetic growth models lies in how metabolic scaling relationships are treated. Classical models, such as those developed by West et al. [47] and more recently by White et al. [49], explicitly assume allometric scaling laws in their theoretical framework. In contrast, our model allows the metabolic scaling exponent and its variability to emerge naturally from the underlying dynamics of stochastic cellular dynamics. Both the West et al. and White et al. models implicitly assume that all metabolic energy is perfectly converted into maintenance, growth, or reproductive output, whereas our model incorporates a thermodynamic constraint by explicitly modeling metabolic inefficiencies, which represents energy that is lost and cannot be used for biomass accumulation, offering a more realistic view of the metabolic costs.

### B. Empirical analysis

Individual-level body mass and basal metabolic rate (*B*) data were compiled from several published databases [21, 42, 50]. We included species in our analysis which had at least 5 individual measurements available for basal metabolic rates and body masses. In total, we analyzed 174 species across six taxa (Table S2 in SM). All metabolic rates were standardized to 25°C, except for those of endothermic species. Empirical species-level allometric exponents 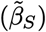 were estimated as the coefficients of log–log regressions of metabolic rate on body mass.

## III. RESULTS

### A. The role of metabolic fluctuations in allometric scaling

We simulated six hypothetical metazoan species across different body sizes (Table S1). For simplicity, we assumed that cell mass (*m*_*c*_), cell metabolism (*B*_*c*_) and the energy required to create new cells (*E*_*c*_) are constant across species and development [48]. We focused on the effect of the stochastic inefficiency coefficient (*γ*) and the baseline fluctuation level (*ϕ*) on the allometric scaling exponents. Without incorporating stochastic inefficiency (i.e. *γ* = 0) or the baseline fluctuation level (i.e. *ϕ* = 0), the allometric scaling exponent *β* between metabolism and body mass is isometric (Fig.2). Adding increasing dissipative term and baseline fluctuation level, the scaling between metabolism and body mass naturally arises, without explicitly imposing a scaling relationship between them (Fig.2a and Fig.S1). The Kleiber’s law exponent (3*/*4) also emerges from the model as a special limiting case, when the linear maintenance term and stochastic jump term of the expected adult metabolic rate (Eq. 9.) are equal (see SM for details).

**Figure 1.**
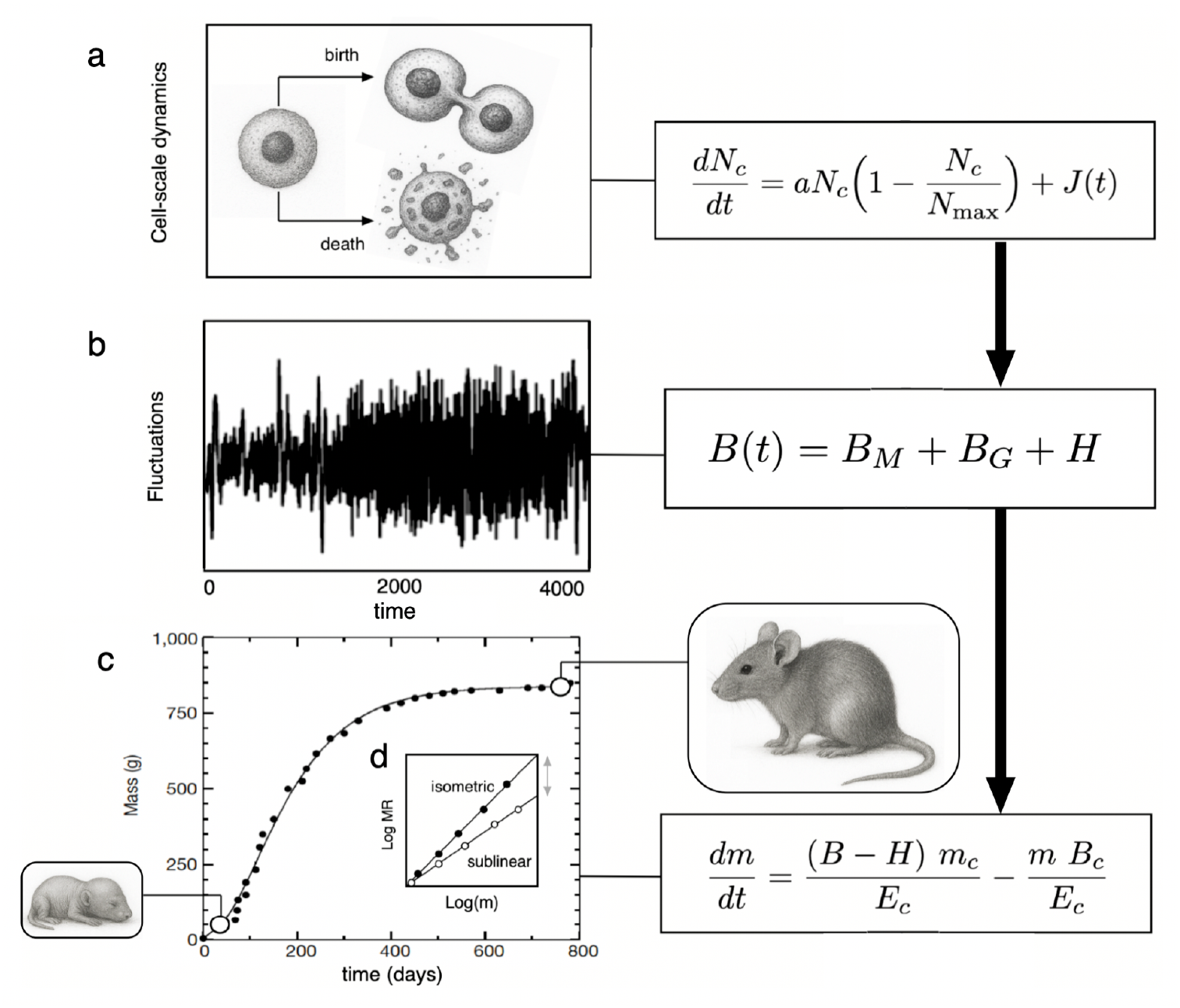
Schematic representation of the stochastic ontogenetic growth model. The stochastic ontogenetic framework is based upon (a) stochastic cellular dynamics, where cells proliferate combined with bursty, stochastic events of cell birth and death. (b) The total basal metabolic rate changes according to cell number change and comprises of cell maintenance, cell growth and metabolic inefficiencies due to stochastic events (plot taken from ref. [26]). (c) Individual body mass changes equals to the difference between metabolic growth and maintenance, which then leads to (d) the scaling relationship between adult mass and metabolism.

**Figure 2.**
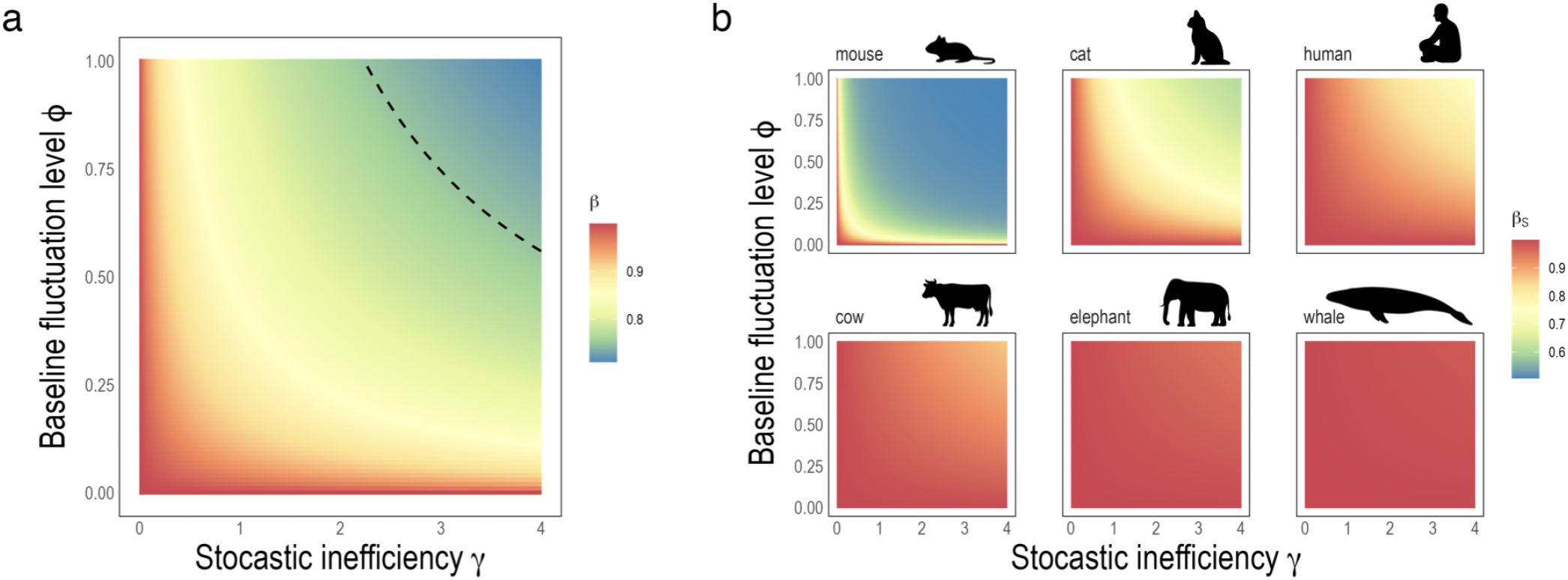
The effect of stochastic inefficiency coefficient (*γ*) and the baseline fluctuation level (*ϕ*) on allometric scaling exponents across species. (a) shows cross-species allometric exponents where the dashed line indicates the canonical value of 3*/*4. (b) shows the variations of species-specific allometric scaling exponents.

The higher the cellular level stochasticity and the sensitivity of the organism to cellular level stochastic processes, the lower the scaling exponent becomes. At low *γ* values, cross-species *β* approaches 1, indicating a near-maximal conversion efficiency of metabolic energy into somatic growth. Species exhibit distinct species specific allometric scaling exponents *β*_*S*_, reflecting differences in developmental timescales and metabolic strategies (Fig.2b). The model results show that the sensitivity to cellular stochastic processes imposes a lower threshold onto the possible allometric scaling exponents. Small-sized animals such as mice can theoretically reach the lowest scaling exponents, whereas the possible scaling exponents for large animals such as whales are constrained in a more narrow range.

Similarly, the model reveals a strong, nonlinear inverse relationship between the stochastic inefficiency, baseline fluctuation level and the overall metabolic fluctuations measured as the standard deviation of the Laplace noise (Fig.S2). These fluctuations in basal metabolic rates have a highly predictable effect on species-specific allometric scaling exponents (Fig.4). Regardless of species, the values of *γ* and *ϕ*, allometric scaling exponents decrease with larger metabolic fluctuations and collapse onto the same line, i.e. that metabolic fluctuations directly predict allometric scaling exponents, providing empirically testable predictions. Overall, these findings suggest that allometric scaling is constrained by individuallevel thermodynamic constraints.

**Figure 3.**
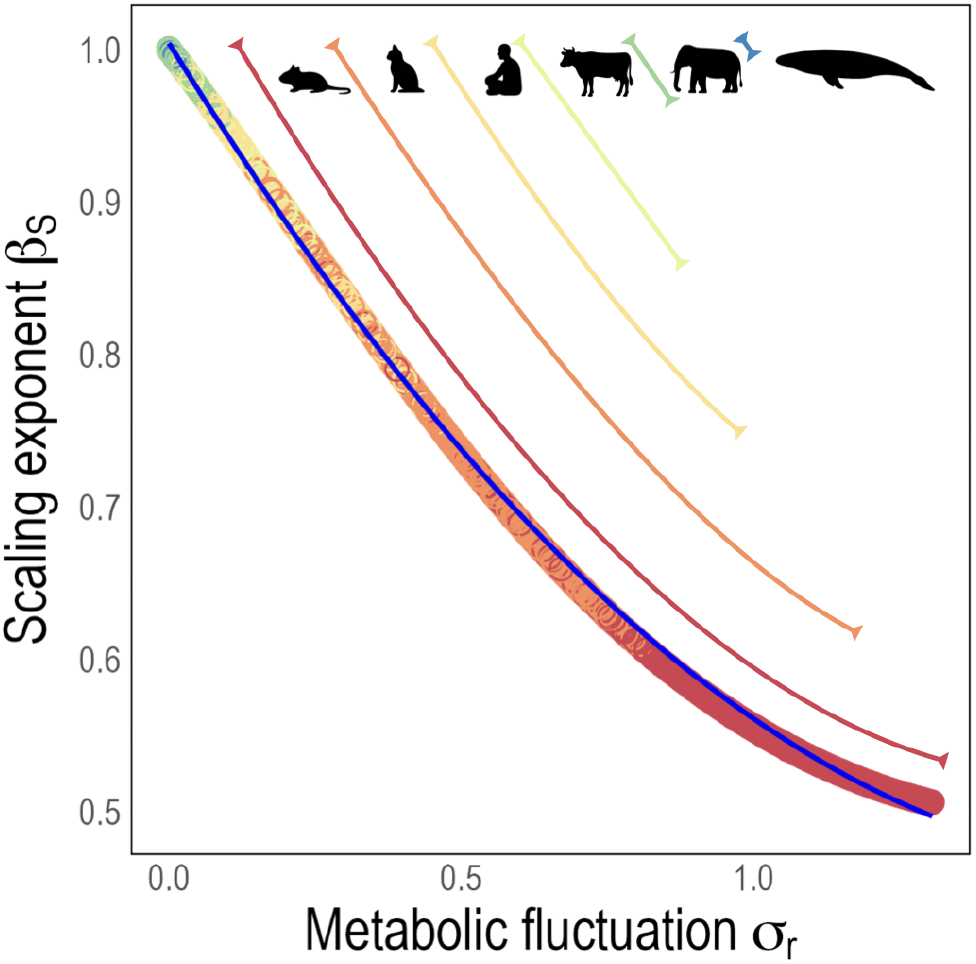
Relationship between metabolic fluctuations and allometric scaling exponent. All *β*_*S*_ exponents across different species and parameter values of *γ* and *ϕ* map onto the same line. The blue line is a fitted quadratic polynomial regression model 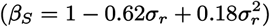. The colored solid lines show how the hypothetical species range on the line.

**Figure 4.**
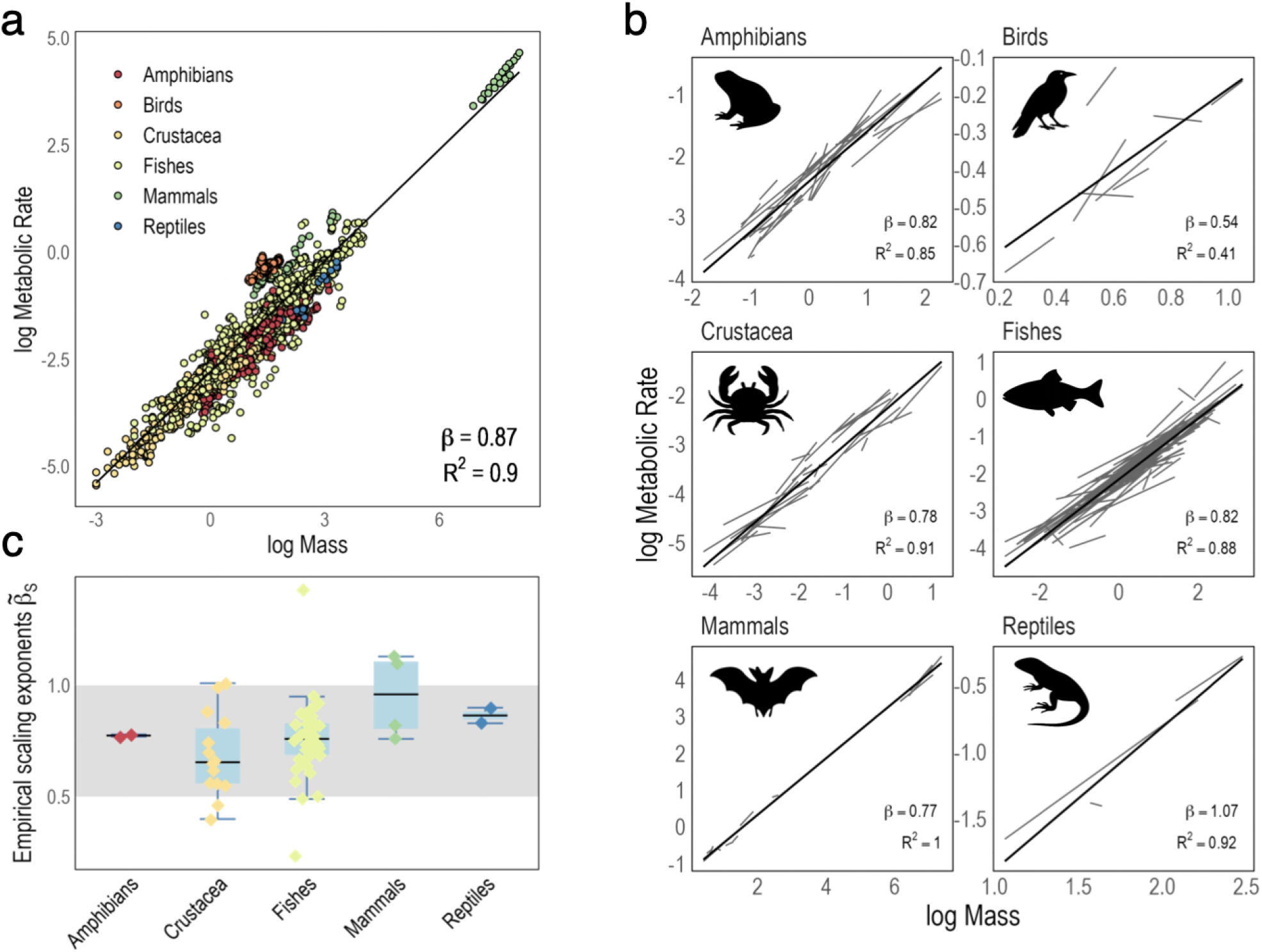
Empirical scaling of metabolic rate across species. (a) Basal metabolic rates vs. body mass calculated for metazoans. The solid black line is a power law fit to the entire dataset. (b) The solid black lines show the power law fits to higher taxa and the grey lines indicate power law regression lines to individual species. (c) The empirical species scaling exponents 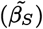 with good model fits (*R*^2^ *>* 0.9) show a close alignment with the range of predicted exponents [0.5, 1].

### B. Diversity of empirical scaling exponents

Metazoans exhibit diverse allometric scaling exponents across taxa. In our empirical sample of species, metazoan body masses scale with metabolic rates with the slope of 0.87, approaching an isometric scaling exponent (Fig.4a). Living things most probably do not show uniform stochastic inefficiencies or baseline fluctuation levels across species. In order to recover an average crossspecies *β* value, we performed a bootstrap analysis on the simulated dataset. In each replicate, we randomly sampled one observation per animal across *γ* and *ϕ* values, then fit a power-law regression of metabolic rate against mass. The resulting bootstrap distribution yielded a mean *β* of 0.82 with a 95% confidence interval of [0.73 – 0.97] similar to the average empirical exponent.

Metazoans grouped into higher taxa, however, show distinct allometric exponents (Fig.4b), which highlights general physiological differences between species groups. Within taxa, species with similar body masses, morphology and anatomy also show highly diverse species-specific scaling exponents pointing towards differing energetic lifestyle and thermoregulatory strategies. However, not all species aligned well with the power law assumption. We then filtered the observed species-level scaling exponents 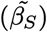 for good model fit (*R*^2^ *>* 0.9), which range closely with the predicted range between [0.5, 1] (Fig.4c). As expected from simulation results, larger endothermic species have close isometric scaling between metabolism and mass, whereas ectothermic species can show a huge variability in their exponents. Deviations from this range can be explained by either inaccuracies in individual measurements and small sample sizes or species-specific physiologies that are not captured by the model.

The observed universality of tent-shaped (Laplace-like) distributions of metabolic fluctuations across species [27] was also recovered in our model. In most cases, a Laplace distribution fits the metabolic fluctuations better than a Gaussian distribution, however, higher fluctuation levels in cell dynamics cause the distribution to become Gaussian (Fig.S3). In the same study, the authors showed power law scaling between the mean absolute deviation of metabolic fluctuations (*σ*_*r*_) and average species body masses as well as average metabolic rates, with the observed scaling exponents of *ν* = −0.241 *±* 0.103 and *δ* = −0.352 *±* 0.072, respectively [27]. Our theoretical model closely recovered these empirical scaling exponents between the average mean absolute deviation of metabolic fluctuations and average body masses *ν* = −0.305*±*0.038 and metabolic rates *δ* = −0.386 *±* 0.025 (see Fig.S4).

## IV. DISCUSSION

The present model provides a mechanistic basis for understanding how scaling between metabolism and body mass emerges from first principles at the level of individual organisms. The fundamental role of metabolic fluctuations in allometric scaling sheds light on the observed diversity of scaling exponents that point toward the importance of lifestyle and thermoregulatory strategies [28] that often override geometric constraints. As shown by Labra et al [27], metabolic fluctuations negatively scale with average metabolic rates. Metabolic fluctuations are expected to be lower in endotherms, due to their regulated homeostasis, which comes with higher metabolic costs and faster tissue turnover. On the other hand, ectotherms’ metabolism tracks ambient conditions relying on behavioral thermoregulation such as shade-seeking or basking [9], which result in lower metabolic rates and slower cellular dynamics and larger fluctuation variability. In this model, stochastic fluctuations in cell number are represented by Laplace noise. The parameter *ϕ* sets the magnitude of these fluctuations, reflecting the intrinsic variability in cell proliferation and loss. The parameters *ϕ* and *γ* translate cell-level stochasticity into metabolic fluctuations. Higher *ϕ* values capture faster tissue turnover and higher per-cell metabolic rates in endotherms, whereas ectotherms are represented by lower *ϕ*, consistent with slower cellular dynamics. The parameter *γ* modulates the energetic consequences of stochastic events, i.e., each deviation in cell number incurs an additional metabolic cost, beyond that required for biosynthesis.

Empirically, the species-level scaling exponents show a huge variability, with only a smaller subset of species fitting statistically to the assumed allometric scaling relationship (Fig.4c). Ectothermic species show larger variability compared to endotherms ranging relatively well within the predicted scaling range. In addition, the negative empirical scaling between metabolic fluctuations and average species body masses and metabolic rates [27] are also recovered in our model, which further confirms the validity of our theoretical predictions (Fig.S4). The universal effect of metabolic fluctuations on allometric scaling exponents across taxa, regardless of their thermal physiology (i.e., *γ* and *ϕ* parameters) or body mass implies that deviation from linear scaling is the direct consequence of stochastic inefficiencies. In other words, the huge variety of observed allometric scaling exponents within and across taxa can be understood through the lens of fluctuations, which departs us from the idea of searching for a single universal exponent.

This work provides a minimal mechanistic framework capturing the primary source of metabolic variability during ontogenetic development. While the model does not explicitly include additional sources of metabolic fluctuations, such as cell size changes, hormonal modulation, mitochondrial variability, behavioral activity, or environmental influences [16, 18, 34, 35], this approach allows us to quantify the contribution of stochastic cell dynamics to metabolic fluctuations. Consequently, the model provides a lower-bound estimate of metabolic fluctuations and a mechanistic baseline for understanding allometric scaling relations, which can be extended in future work to incorporate additional physiological and environmental factors. Furthermore, this model is most suitable for modeling multicellular animals. While thermogenesis is universal among animals, with some rare exceptions, plants are generally not thermogenic [38]. Thus the energetic impact of stochastic fluctuations in cell number is way smaller in plants compared to animals and primarily arise during active growth in meristematic tissue rather than throughout the whole organism.

The broader relevance of this framework for other complex system scaling laws lies in linking microscopic fluctuations to macroscopic patterns. The model reveals that metabolic scaling is a consequence of thermodynamic inefficiencies bridging the gap between physiological mechanisms and larger ecological patterns. Stochastic processes rooted in random bursty events of cellular dynamics introduce intrinsic variability that propagates from cellular to organismal level, shaping emergent thermodynamic constraints. Overall, this work offers a unified framework to understand metabolic scaling laws and to interpret the diversity of observed scaling relationships across complex organisms.

## ACKNOWLEDGMENTS

RS acknowledges support from an AGAUR 2021 SGR 0075 grant, a Grant PID2023-152129NB-I00 funded by MICIU/AEI/10.13039/501100011033 and the Santa Fe Institute.

## AUTHOR CONTRIBUTIONS

Conceptualization: A.T.; Methodology: A.T.; Formal Analysis: A.T.; Writing – Original Draft: A.T.; Validation: A.T., W.M., F.A.N.S., R.S.; Writing – Review & Editing: A.T., W.M., F.A.N.S., R.S.

## Supplementary Materials

## S1 Materials and Methods

### Stochastic Ontogenetic Growth Model

This model simulates ontogenetic growth using the principle of energy conservation at the cellular level, incorporating stochasticity in cell dynamics [**?**]. The organism’s mass changes over time based on cellular-level metabolic and growth dynamics [**?**]. The input parameters are the mean maximum number of cells 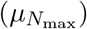, the variation of the maximum number of cells 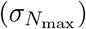 per species, the baseline fluctuation level of cell dynamics (*ϕ*) and the stochastic inefficiency (*γ*).

### S1.1 Model Inputs

#### Fixed Model Parameters

- *m*_*c*_: Mass of a single cell (kg)
- *B*_*c*_: Basal metabolic rate per cell (W = J/s)
- *E*_*c*_: Average energy required to create a new cell (J)

#### Tuning Model Parameters

- *γ*: Stochastic inefficiency coefficient (J/cell)
- *ϕ*: Baseline fluctuation level

#### Species-specific Model Parameters

- *a*: Cellular growth rate (cell^−1^s^−1^) calculated as 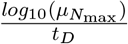
- *t*_*D*_: development time
- 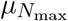: Log-normal mean parameter for *N*_*max*_ (maximum number of cells)
- 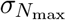: Log-normal noise parameter for *N*_*max*_ (maximum number of cells)

#### Time-dependent Variables

- *m*(*t*): Total organism mass at time *t* (kg)
- *N*_*c*_(*t*): Number of cells at time *t*
- *B*(*t*): Total metabolic rate at time *t* (W)

#### Individual-specific Parameters

- In order to add individual-level noise to masses, *N*_max_ (the maximum number of cells) is sampled from a log-normal distribution for each individual:

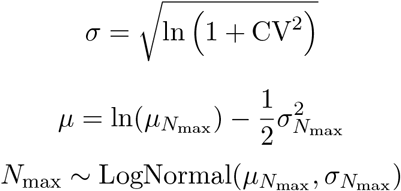

### S1.2 Model Equations

#### Stochastic Cell dynamics with Jump process

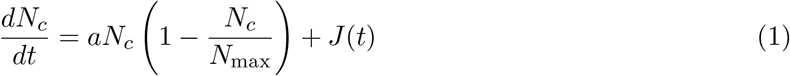

J(t) is approximated via symmetric exponential jumps [**?**]:

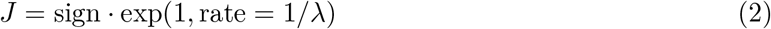

Using *n* = *N*_*c*_*/N*_max_, let us assume that each of the *N*_*c*_ cells can, with probability *π*(*n*), produce a stochastic burst during a short time window Δ*t*. Each burst has:

- An assigned random sign *S*_*i*_ ∈ {+1, −1}, with equal probability,
- Exponential amplitude *E*_*i*_ ∼ exp(1*/λ*_0_), i.i.d. with mean *λ*_0_.

Let *K*, which follows a binomial distribution, that is, *K* ∼ *B*(*N*_*c*_, *π*(*n*)), be the number of cells. Then the total fluctuation can be written as

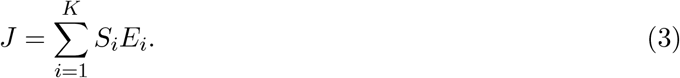

Since 𝔼 [*S*_*i*_*E*_*i*_] = 0 and 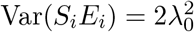, the variance of *J* becomes

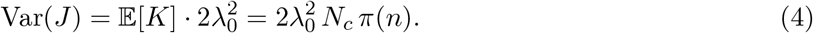

We now approximate this by a single symmetric Laplace-distributed jump:

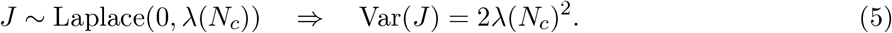

Matching variances gives:

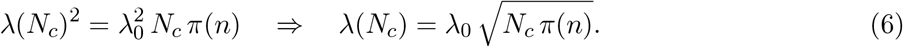

To define *π*(*n*), we introduce a developmental modulation of variability. The function *π*(*n*) represents the normalized activity or burst probability per cell, and we require:

- *π*(0) = 1: maximal variability at early development (unregulated growth),
- *π*(1) = *ϕ*^2^: residual noise at confluence (fully developed system),
- *π*(*n*) is monotonic decreasing in *n*, capturing developmental canalization.

A minimal smooth function that satisfies these criteria is:

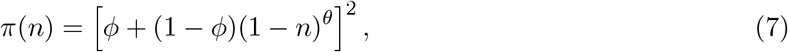

where:

- *ϕ* controls the residual variability at *n* = 1,
- *θ >* 0 is a steepness parameter that tunes how rapidly variability declines with development.

The form (1 − *n*)^*θ*^ reflects resource/niche availability or degrees of freedom for cell fate. The exponent *θ* modulates this transition. Specifically, we have:

- *θ* = 1 indicates a linear decay in variability,
- *θ <* 1 implies slow decay, i.e. variability persists longer into development,
- *θ >* 1 implies rapid decay i.e. variability drops sharply as *N*_*c*_ increases.

This allows the model to capture a range of developmental noise dynamics, from gradual stabilization to sharp canalization.

Plugging this into the expression for *λ*(*N*_*c*_) and setting *λ*_0_ = 1 (by rescaling units) yields:

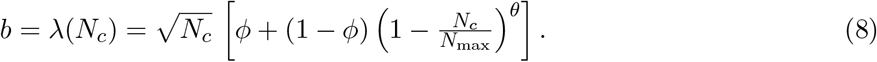

This function grows as 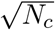 for small *N*_*c*_, matching the expected demographic scaling, and decays toward 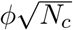 as *N*_*c*_ → *N*_max_, capturing reduced stochasticity as development proceeds.

In our simulations, we adopted a discrete time step of Δ*t* = 0.1 h to numerically integrate the stochastic ontogenetic growth equations. This choice reflects a compromise between numerical stability and computational efficiency. Given that the characteristic growth rate of the system is on the order of *a* ∼ 10^−2^ − 10^−4^ h^−1^, the deterministic term changes by less than 10^−3^ − 10^−5^ per iteration, ensuring that the Euler integration remains well within the stability regime (*a*Δ*t* ≪ 1). The stochastic Laplace scale term (*b*) was scaled by 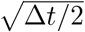 to preserve the correct variance of fluctuations across time resolutions, following the standard diffusion approximation [**?**]. The chosen Δ*t* = 0.1 thus provides smooth trajectories, accurate fluctuation statistics, and consistent steady-state behavior while maintaining computational tractability.

#### Total Metabolic Rate and body mass dynamics

We describe a decomposition of total metabolic rate into three terms:

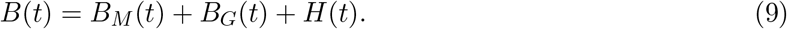

The maintenance component is proportional to the number of cells:

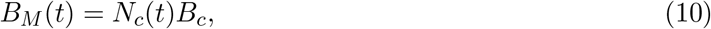

where *B*_*c*_ is the maintenance power expenditure per cell. The growth or turnover component accounts for the energetic cost of producing or removing cells:

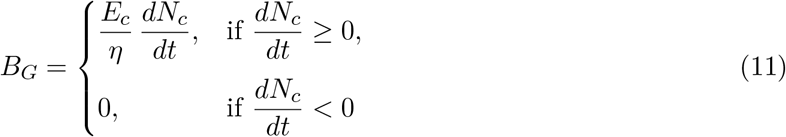

where *E*_*c*_ is the energy required for birth of cells, and *η* is the growth efficiency which fixed to 0.7.

The metabolic inefficiency component reflects metabolic bursts [**?, ?**] and is modeled as proportional to the magnitude of the jump term:

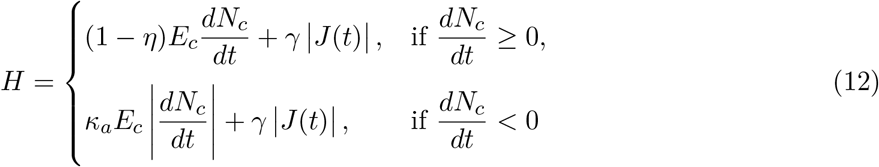

where *J*(*t*) is a stochastic term and *γ* is a stochastic inefficiency coefficient that converts the stochastic burst into energy dissipation.

The cell population evolves according to a stochastic logistic model already described, namely:

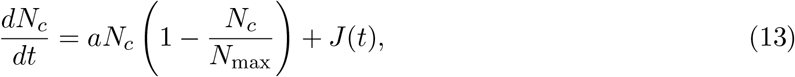

Since *J*(*t*) ∼ Laplace(0, *λ*(*N*_*c*_)), the expected magnitude of metabolic inefficiency is:

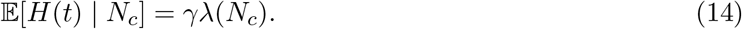

Let *m*(*t*) denote total body mass and *m*_*c*_ the average cell mass. The rate of change of mass is driven by net energy available for biomass production, after subtracting maintenance and metabolic inefficiency. Converting energy into mass at a cost *E*_*c*_ per cell, we obtain:

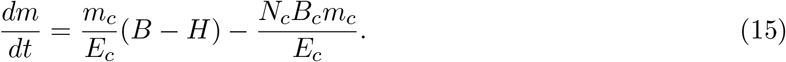

Substituting *N*_*c*_ ≈ *m/m*_*c*_ into the maintenance term gives:

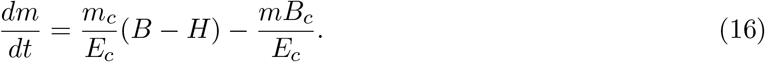

This final form describes the dynamics of body mass based on energy input, metabolic noise, and maintenance cost.

### S1.3 Model Outputs

#### Mean absolute deviation of metabolic fluctuations

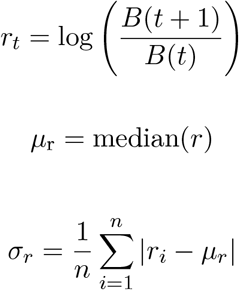

#### Allometric scaling exponent

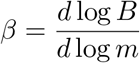

using data where *m* is the mass at maturation and *B* is the metabolic fluctuation in mature state.

### S1.4 Model parametrization

**Table S1.** Model parameters across different body sizes. Data sources: *B*_*c*_ from ref.[1], *m*_*c*_ from ref.[2], *E*_*c*_ from ref.[3].

| Animal | Mass (kg) | Dev. Time | $N_{max}$ | $m_c$ (kg) | $B_c$ (J/hr) | $E_c$ (J) | $CV_{N_{max}}$ |
| --- | --- | --- | --- | --- | --- | --- | --- |
| Mouse | $\sim 0.01$ | $\sim 30$ days | 1e10 | 1e-12 | 1.08e-07 | 3.7e-7 | 0.2 |
| Cat | $\sim 5$ | $\sim 1$ year | 5e12 | 1e-12 | 1.08e-07 | 3.7e-7 | 0.2 |
| Human | $\sim 70$ | $\sim 18$ years | 7e13 | 1e-12 | 1.08e-07 | 3.7e-7 | 0.2 |
| Cow | $\sim 500$ | $\sim 4$ years | 5e14 | 1e-12 | 1.08e-07 | 3.7e-7 | 0.2 |
| Elephant | $\sim 5,000$ | $\sim 25$ years | 5e15 | 1e-12 | 1.08e-07 | 3.7e-7 | 0.2 |
| Whale | $\sim 50,000$ | $\sim 30$ years | 5e16 | 1e-12 | 1.08e-07 | 3.7e-7 | 0.2 |

### S1.5 Estimation of Metabolic Fluctuations *σ*_*r*_

Metabolic fluctuations in the adult phase are quantified from

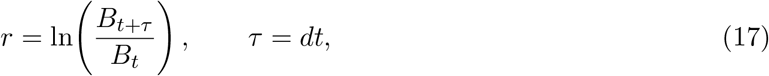

where the only difference between *B*_*t*_ and *B*_*t*+*τ*_ arises from a stochastic jump in cell number. In the simulation the jump term is drawn from a Laplace distribution

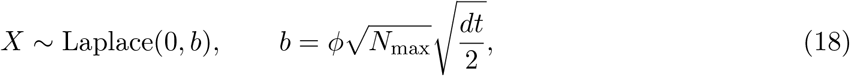

and enters the metabolic balance

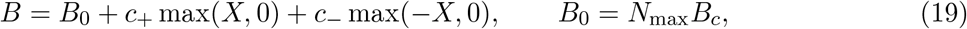

With

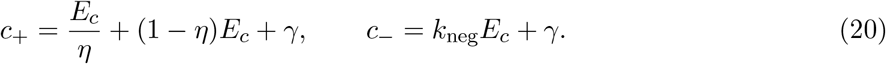

The fluctuation amplitude reported in all results is the mean absolute deviation

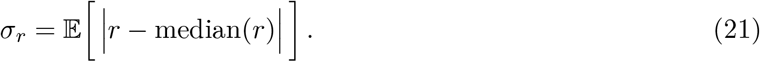

To avoid bias across the whole parameter regimes, we estimated *σ*_*r*_ numerically by Monte Carlo sampling. For each parameter set we draw *X*_1_, 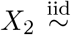 Laplace(0, *b*), to construct *B*_*t*_ and *Bt*+*τ* and compute the sample MAD of over 2 *r* × 10 replicates. This procedure yields unbiased estimates that match the simulated time-series statistics for all species sizes.

### S1.6 Calculation of the Metabolic Scaling exponent *β*_*S*_

At adult steady state (*N*_*c*_ → *N*_max_) the expected value becomes

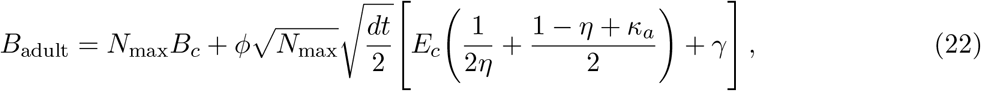

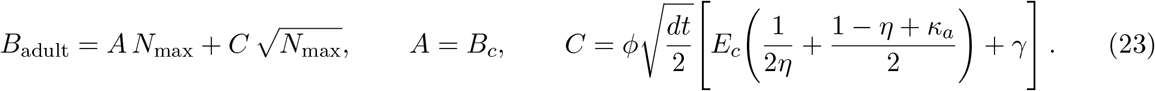

Similarly, the expected adult mass

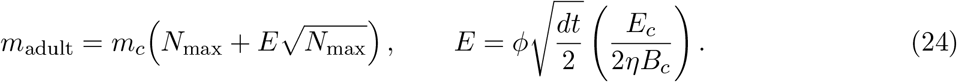

The metabolic scaling exponent *β*_*S*_ is defined as

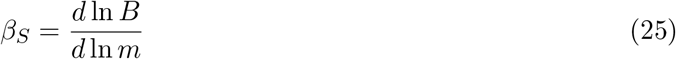

Substituting the adult relations gives

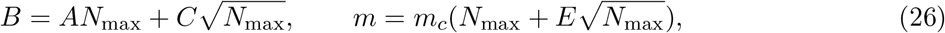

Differentiate with respect to *N*_max_:

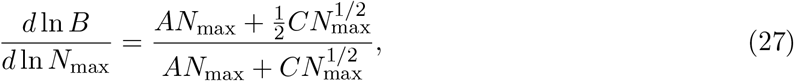

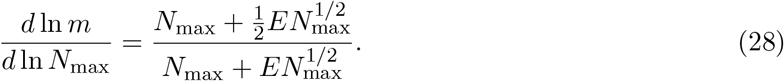

Hence

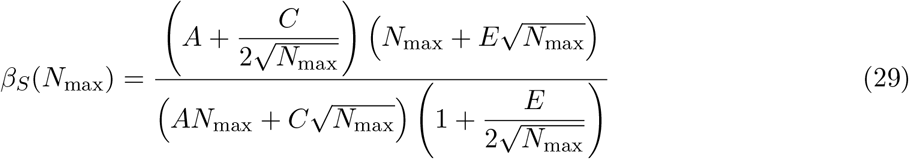

For large *N*_max_ (large species),

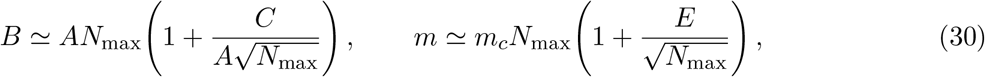

so

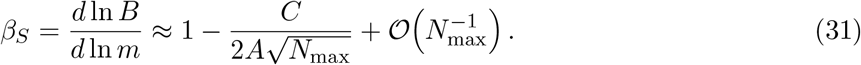

Thus *β*_*S*_ decreases slightly below 1 as body size grows *β*_*S*_ *<* 1. The maintenance term scales linearly with cell number: *B*_*M*_ ∝ *N*_*c*_. The stochastic term scales sublinearly: 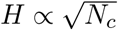 (central-limit behavior of fluctuations). Their sum produces an emergent power law between total metabolism and body mass:

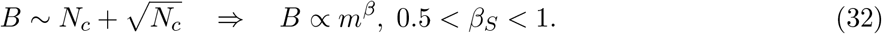

Hence, the sublinear metabolic scaling arises *naturally* from combining deterministic and stochastic energetic components in the growth model.

### S1.7 Derivation of the 3*/*4 Exponent

Let 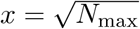 and write the metabolic scaling exponent as

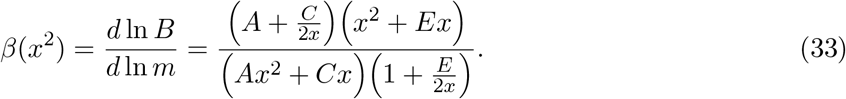

We now evaluate *β* under the special balance condition [**?, ?**]

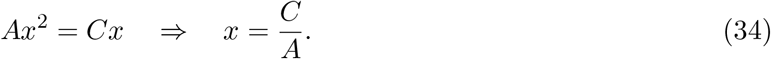

Then, the individual factors simplify:

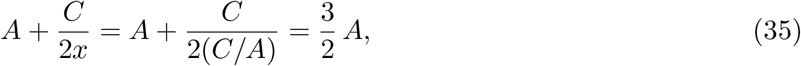

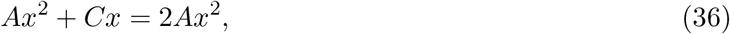

Then we obtain

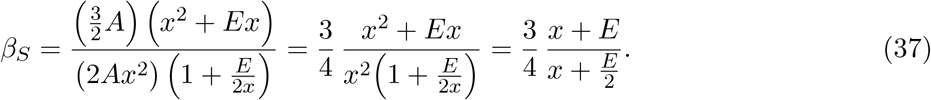

Two limiting cases are immediate:

1. No mass correction (*E* = 0).

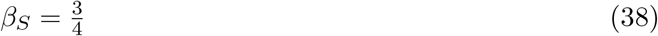
2. Finite *E* but large body size 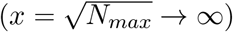.

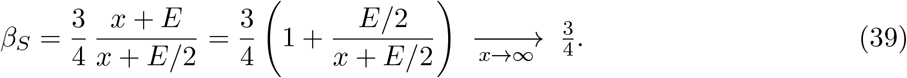

Thus the model predicts the canonical 3*/*4 metabolic exponent in the efficient limit, i.e., if the linear maintenance *AN*_*max*_ and stochastic jump contribution 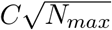 are equal [**?, ?**], and approaches ¾ asymptotically for large species [**?, ?**] .

### S1.8 Microscopic birth–death–heat model underlying the stochastic ontogenetic framework

#### Notations

- *X*(*t*) (integer) – number of cells at time *t*.
- *H*(*t*) (continuous) – bookkeeping variable that counts cumulative heat quanta released by cellular events and relaxes to the environment at rate *γ. Units*: arbitrary “heat units”.
- *k*_*b*_, *k*_*d*_, *k*_*c*_ – birth, death, and competition rate constants (time^−1^). Assumed constants throughout ontogeny.
- *ν*_*b*_, *ν*_*d*_, *ν*_*c*_ – heat yield per event (*ν* ≥ 0, dimension = heat).
- Ω – system-size parameter (e.g. body volume).
- 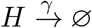 – exponential heat leak with decay rate *γ* (time^−1^).

#### Reaction scheme

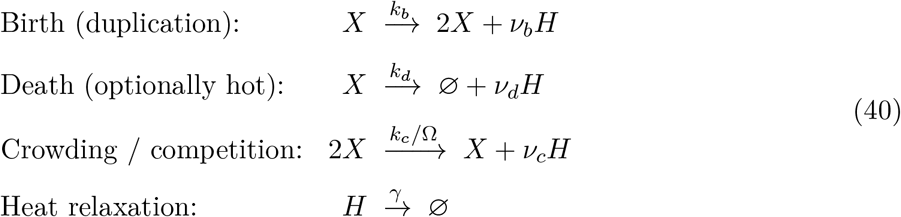

The stochastic ontogenetic model aggregates heat loss into the metabolic inefficiency term *H* in Eq. (12). Making *H* explicit clarifies *where* heat comes from and *why* its fluctuations are Laplace-distributed.

#### From reactions to the chemical master equation

For a small time step Δ*t* the probability of exactly one birth is *k X* Δ*t* + 𝒪 (Δ*t*^2^), one death is *k*_*d*_*X* Δ*t* + 𝒪 (Δ*t*^2^), etc. Let *P* (*n, h, t*) denote Pr [*X*(*t*) = *n, H*(*t*) = *h*] . Enumeration of all one-step events yields:

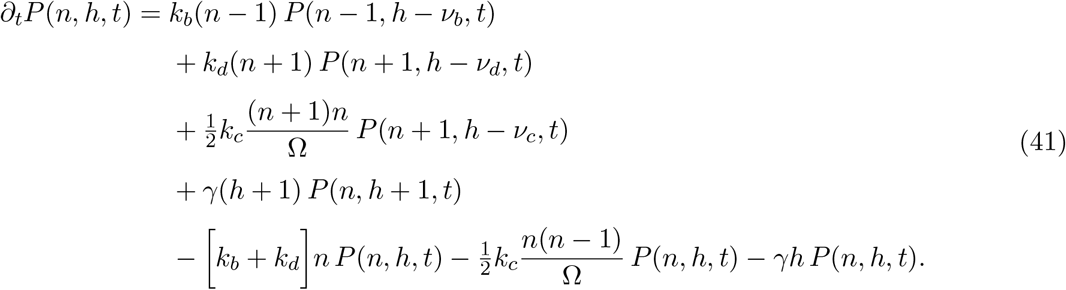

This is the master equation associated to the above reaction network.

#### Mean–field (moment) equations

After multiplying Eq. (21) by *n* and summing over all *n, h*, we obtain the equation for the expectation value for the number of cells *X*

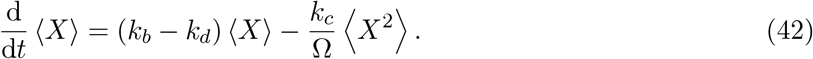

And similarly for the heat *H*:

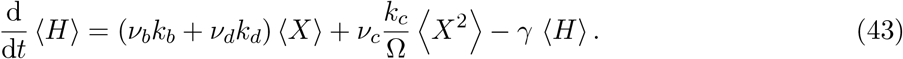

Equations (22)–(23) are *exact* but they involve the second moment ⟨*X*^2^⟩. For a large population, the relative fluctuation 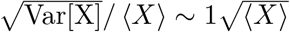 is small, hence ⟨*X*⟩^2^ ≃ ⟨*X*⟩^2^ . Set *N*_*c*_(*t*) := ⟨*X*(*t*)⟩. Then

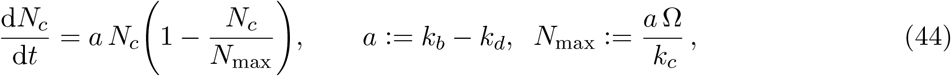

which is the *deterministic logistic* growth equation.

#### Derivation of the jump term

Write *X*(*t*) = *N*_*c*_(*t*) + *δX*(*t*) with ⟨*δX*⟩ = 0. Standard system-size expansion (Kramers–Moyal) [**?**] gives the linear stochastic differential equation

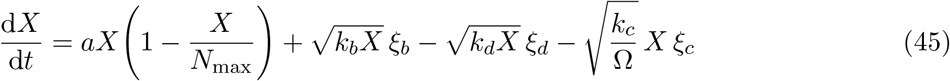

where each *ξ*() is Gaussian white noise (⟨*ξ*(*t*)*ξ*(*t*^*′*^)⟩ = *δ*(*t*−*t*^*′*^)) obtained by centering the corresponding Poisson process. For a short time window Δ*t*, the birth noise increments a random amount Δ*B* ∼ Exp(*λ*_*b*_ = *k*_*b*_*X*), the death noise increments Δ*D* ∼ Exp(*λ*_*d*_ = *k*_*d*_*X*), and the two are *independent*. The net jump Δ*J* := Δ*B* − Δ*D* therefore has the probability density

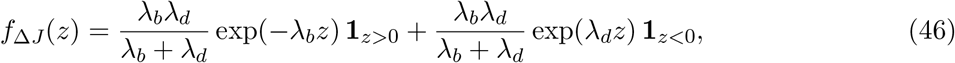

which is the **Laplace distribution** with scale *b* = 1*/*(*λ*_*b*_ + *λ*_*d*_) = 1*/*((*k*_*b*_ + *k*_*d*_)*X*). Hence the absolute jump size |Δ*J*| is *exponential*.

#### Heat bookkeeping SDE

Insert *X* ≈ *N*_*c*_ into Eq. (23) and add the fluctuation terms:

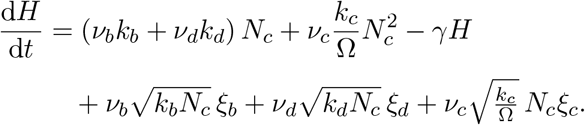

The stochastic part is again Laplace for the same reason as above (sum and difference of exponentials).

## S2 Supplementary Figures

**Figure S1.**
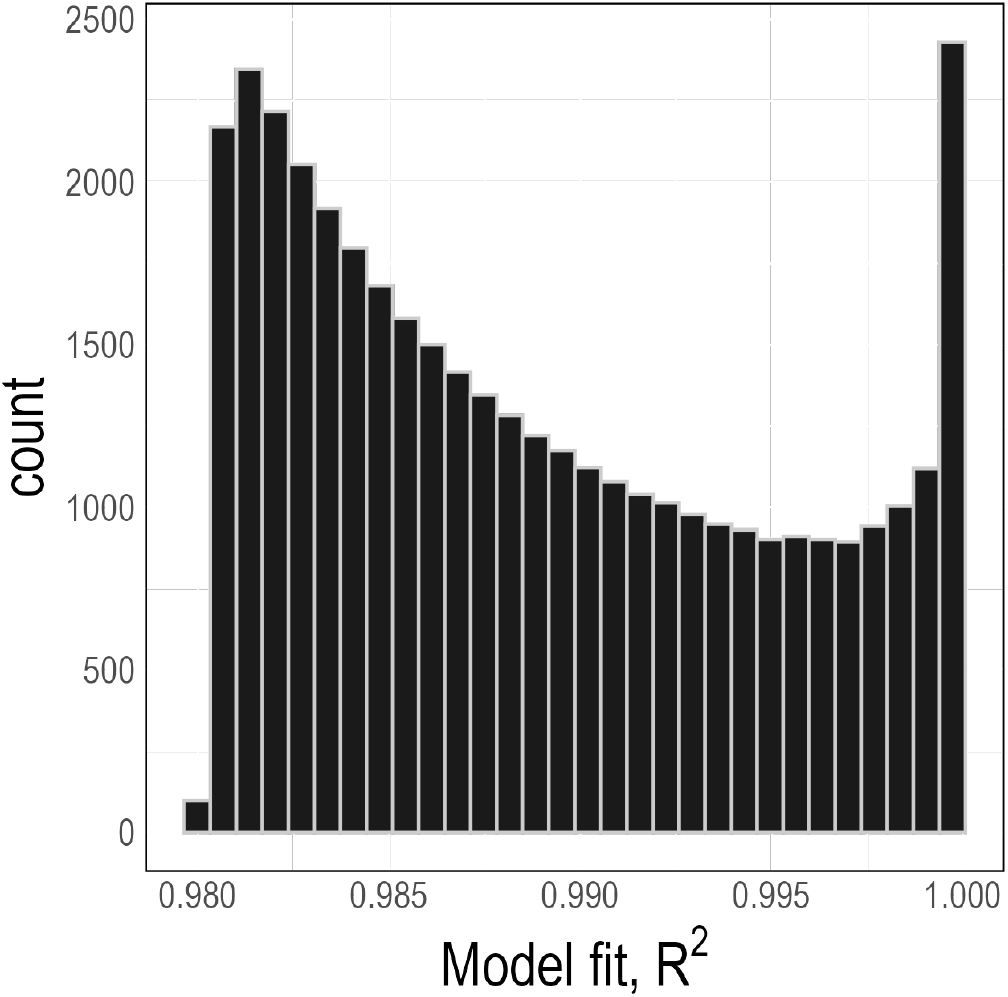
Log-log regression model fits. *R*^2^ values extracted from fitted log-log regression models to simulated values across the six hypothetical species to obtain the *β* across species for Figure 2a.

**Figure S2.**
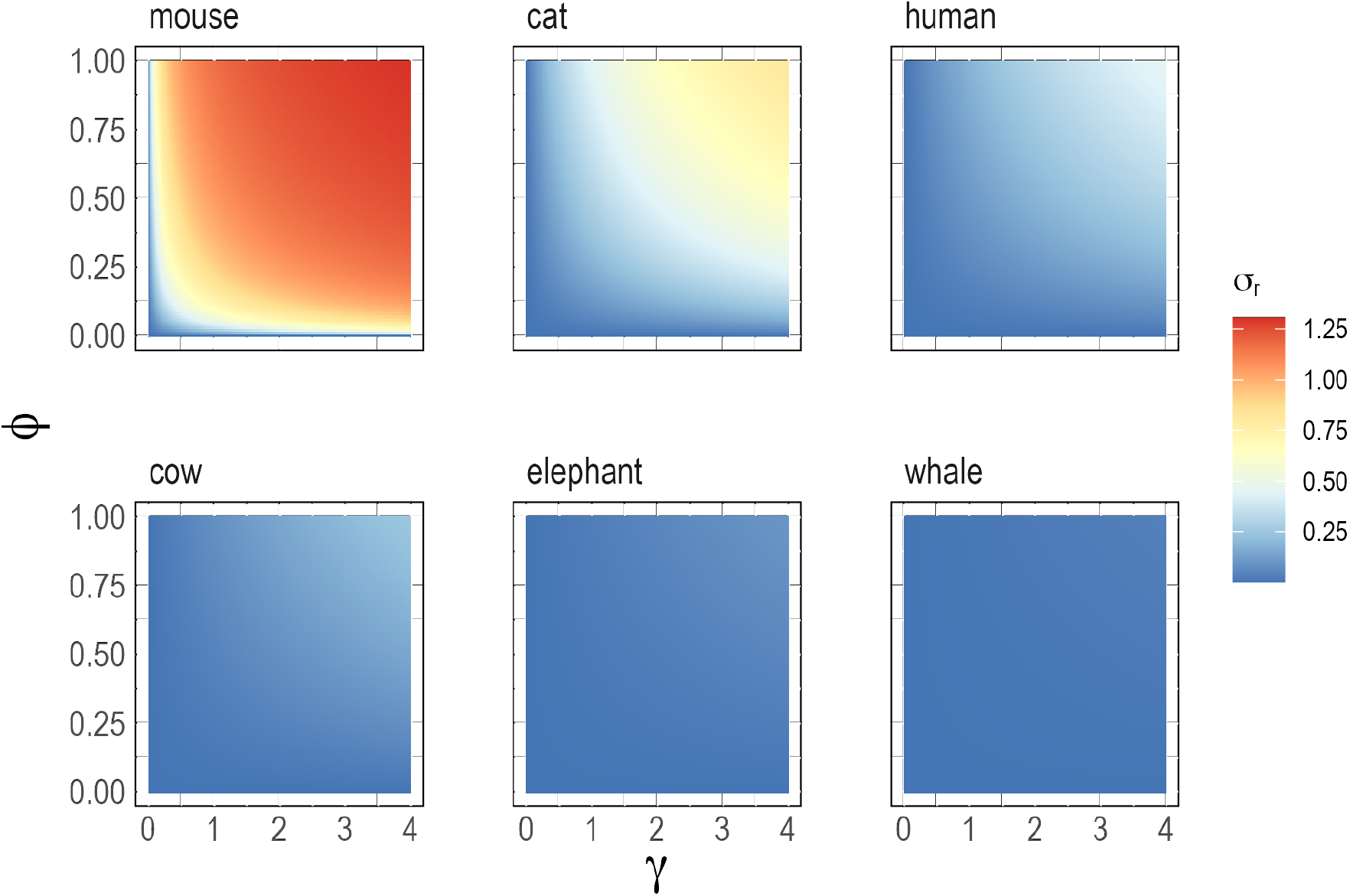
The effect of stochastic inefficiency coefficient (*γ*) and the baseline fluctuation level (*ϕ*) on metabolic fluctuations (*σ*_*r*_) across species. Each point represents the values of (a) ontogenetic and (b) species-specific allometric scaling exponents and metabolic fluctuations measured as the mean absolute deviations in fluctuations of metabolic rates at maturity. Colors denote different values of the fluctuation cost parameter *γ*.

**Figure S3.**
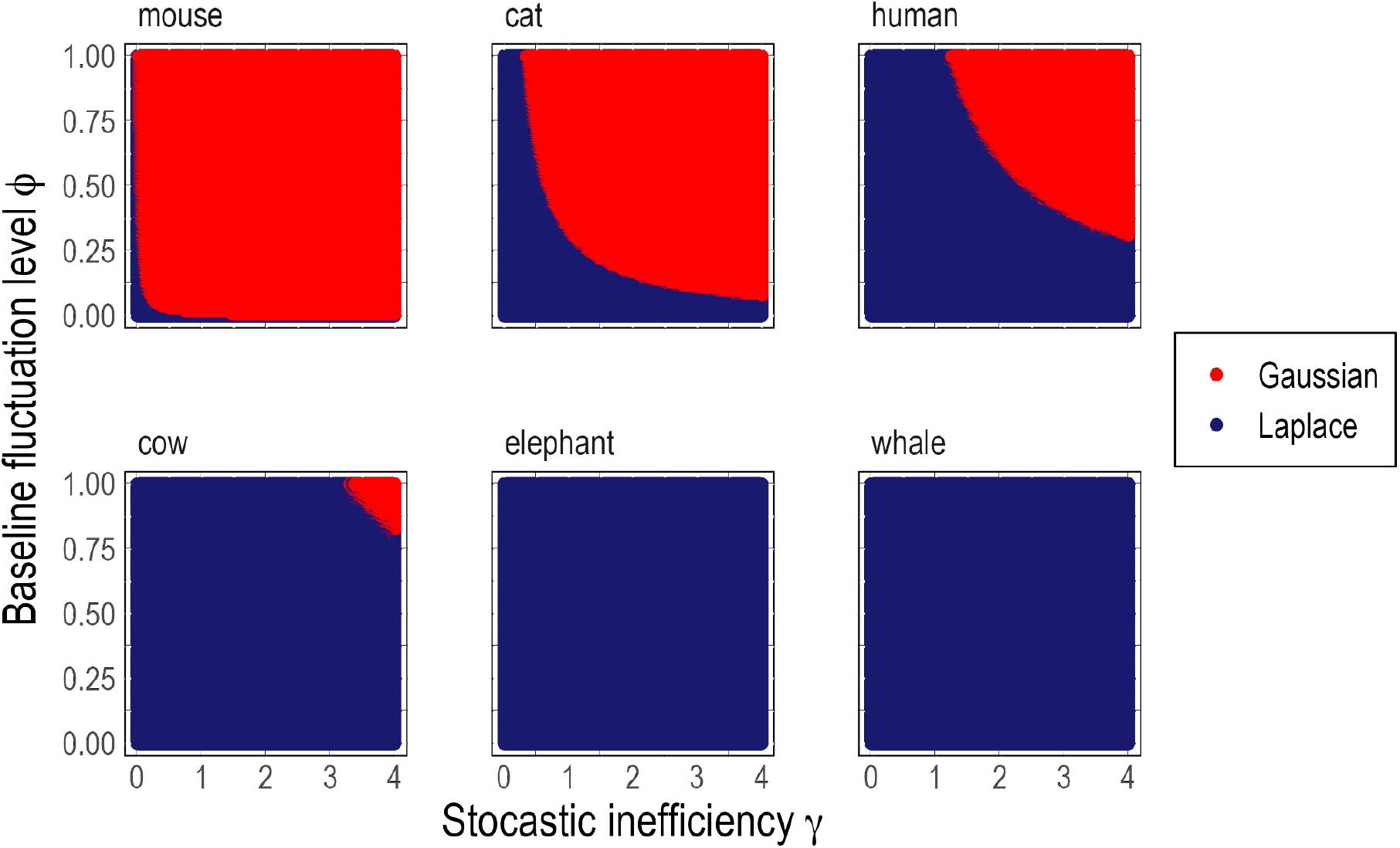
Best fit distributions to metabolic rate fluctuations. Both Gaussian and Laplace distributions were fitted to *σ*_*r*_ time series of simulated data.

**Figure S4.**
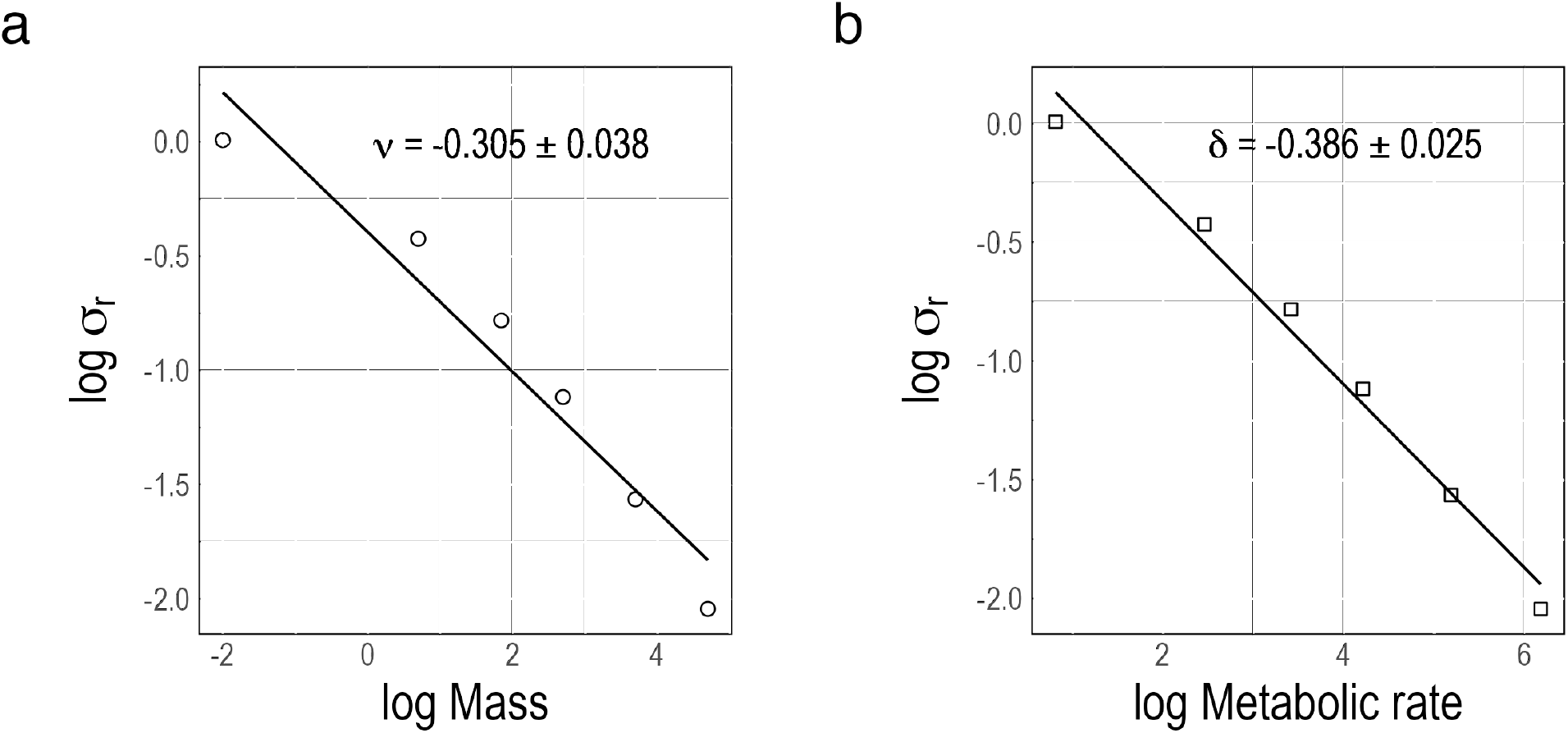
Scaling of metabolic rate fluctuations. Theoretical scaling exponents between the average standard deviation of metabolic fluctuations and (a) average body masses and (b) metabolic rates.

## S3 Data

The body mass and *B* columns contain the averaged values across individuals.

**Table S2.**
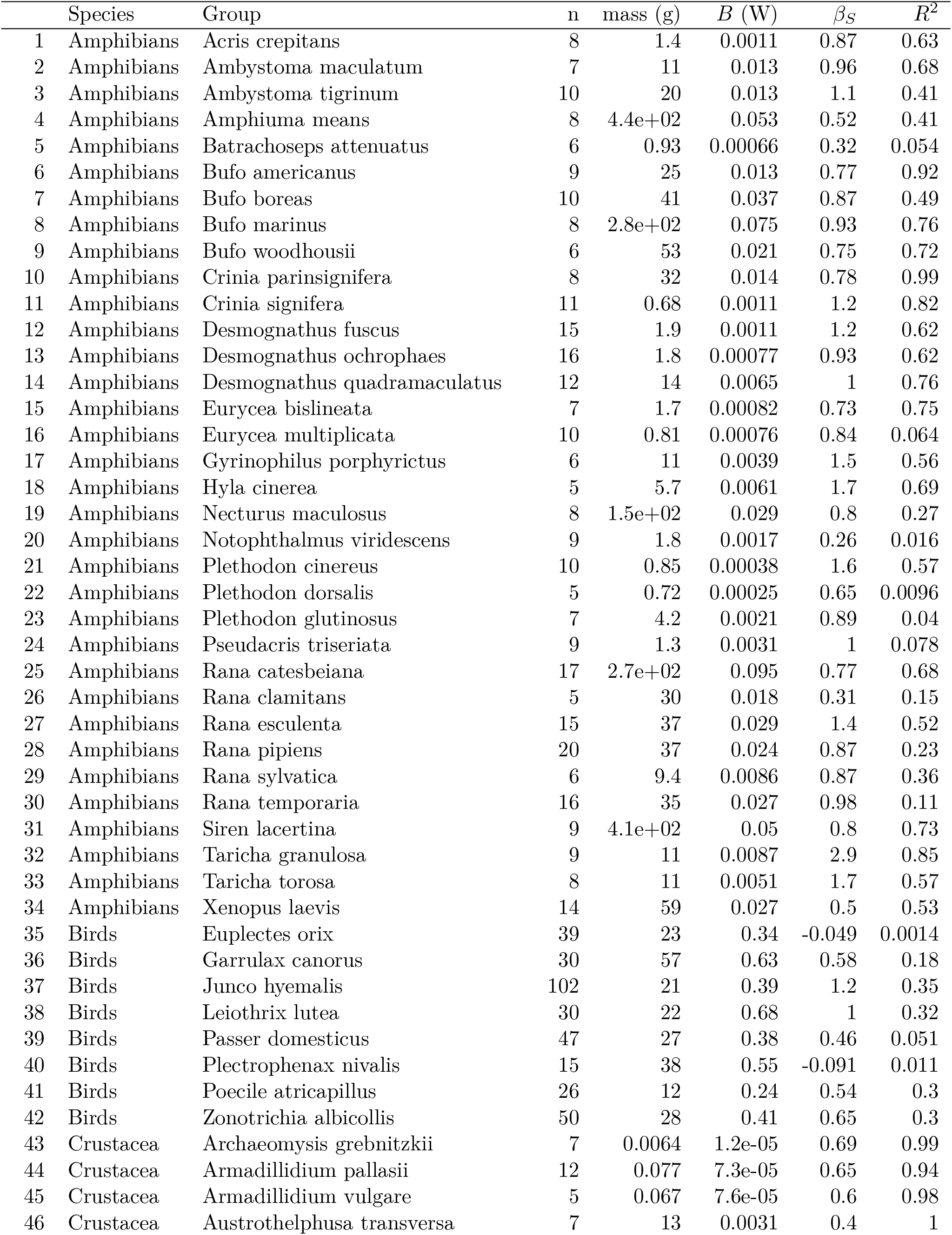
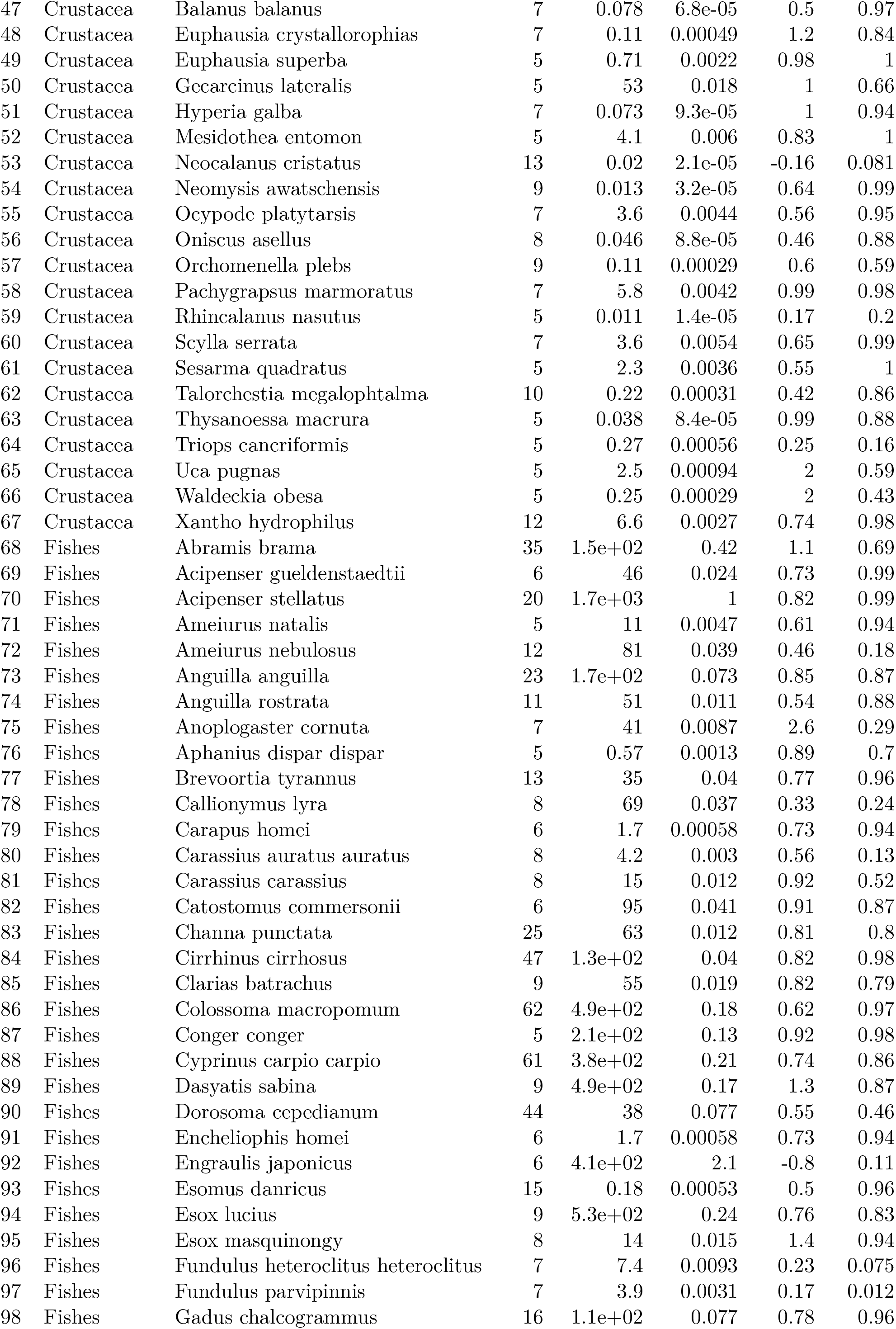
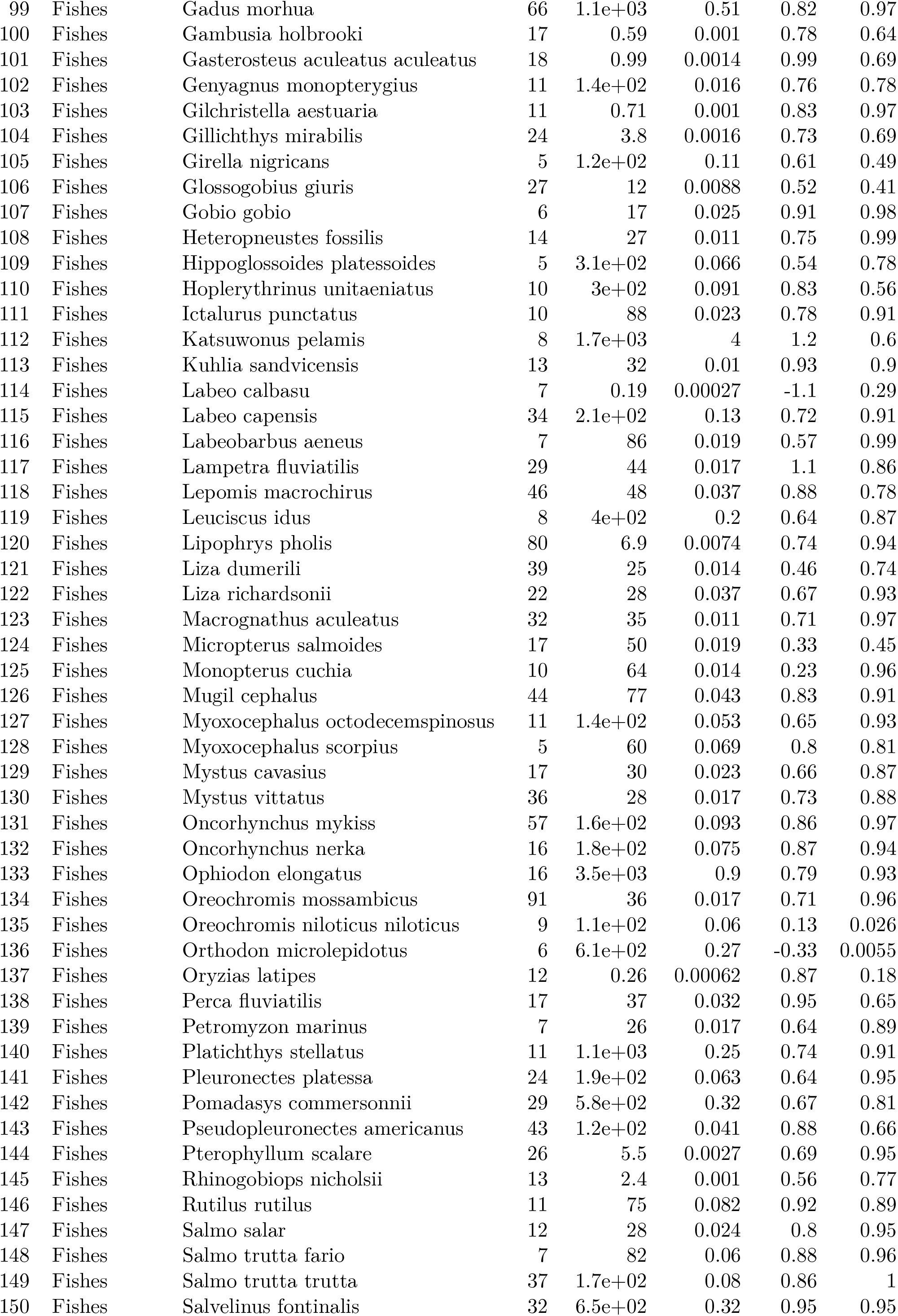
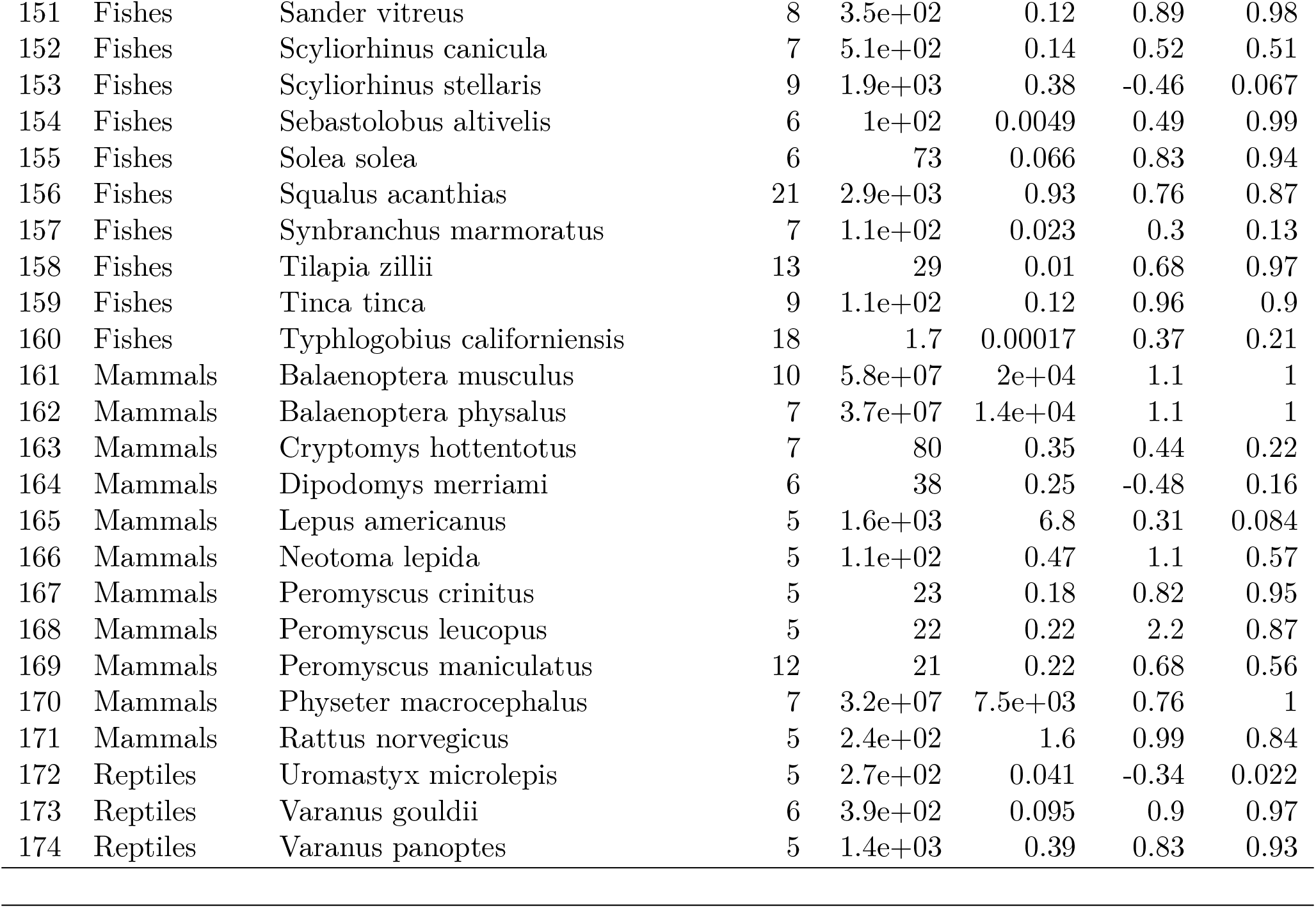
Summary of empirical data.

## REFERENCES

[1] Agutter, P. S. and Wheatley, D. N. (2004). Metabolic scaling: consensus or controversy? Theoretical Biology & Medical Modelling, 1:13.

[2] Artime, O. and De Domenico, M. (2022). From the origin of life to pandemics: emergent phenomena in complex systems. Philosophical Transactions of the Royal Society A: Mathematical, Physical and Engineering Sciences, 380(2227):20200410.

[3] Ballesteros, F. J., Martinez, V. J., Luque, B., Lacasa, L., Valor, E., and Moya, A. (2018). On the thermodynamic origin of metabolic scaling. Scientific Reports, 8(1):1448.

[4] Banavar, J. R., Moses, M. E., Brown, J. H., Damuth, J., Rinaldo, A., Sibly, R. M., and Maritan, A. (2010). A general basis for quarter-power scaling in animals. Proceedings of the National Academy of Sciences of the United States of America, 107(36):15816–15820.

[5] Barabasi, A.-L. and Oltvai, Z. N. (2004). Network biology: understanding the cell’s functional organization. Nature reviews genetics, 5(2):101–113.

[6] Brown, J. H., Gillooly, J. F., Allen, A. P., Savage, V. M., and West, G. B. (2004). Toward a Metabolic Theory of Ecology. Ecology, 85(7):1771–1789.

[7] Brown, J. H. and West, G. B. (2000). Scaling in biology. Oxford University Press.

[8] Christensen, K. and Moloney, N. R. (2005). Complexity and criticality, volume 1. Imperial College Press.

[9] Clarke, A. (2017). Principles of Thermal Ecology: Temperature, Energy, and Life. Oxford University Press.

[10] Clarke, A., Rothery, P., and Isaac, N. J. B. (2010). Scaling of basal metabolic rate with body mass and temperature in mammals. Journal of Animal Ecology, 79(3):610–619.

[11] DeLong, J. P., Okie, J. G., Moses, M. E., Sibly, R. M., and Brown, J. H. (2010). Shifts in metabolic scaling, production, and efficiency across major evolutionary transitions of life. Proceedings of the National Academy of Sciences, 107(29):12941–12945.

[12] Dodds, P. S., Rothman, D. H., and Weitz, J. S. (2001). Re-examination of the “3/4-law” of Metabolism. Journal of Theoretical Biology, 209(1):9–27.

[13] Economos, A. C. (1979). Gravity, metabolic rate and body size of mammals. The Physiologist, 22(6):S71–72.

[14] Giometto, A., Altermatt, F., Carrara, F., Maritan, A., and Rinaldo, A. (2013). Scaling body size fluctuations. Proceedings of the National Academy of Sciences, 110(12):4646–4650.

[15] Gisiger, T. (2001). Scale invariance in biology: coincidence or footprint of a universal mechanism? Biological Reviews, 76(2):161–209.

[16] Glazier, D. S. (2009). Activity affects intraspecific body-size scaling of metabolic rate in ectothermic animals. Journal of Comparative Physiology. B, Biochemical, Systemic, and Environmental Physiology, 179(7):821–828.

[17] Glazier, D. S. (2010). A unifying explanation for diverse metabolic scaling in animals and plants. Biological Reviews of the Cambridge Philosophical Society, 85(1):111– 138.

[18] Glazier, D. S. (2022). How Metabolic Rate Relates to Cell Size. Biology, 11(8):1106.

[19] Grilli, J. (2020). Macroecological laws describe variation and diversity in microbial communities. Nature Communications, 11(1):4743.

[20] Heusner, A. A. (1982). Energy metabolism and body size. I. Is the 0.75 mass exponent of Kleiber’s equation a statistical artifact? Respiration Physiology, 48(1):1–12.

[21] Hoehler, T. M., Mankel, D. J., Girguis, P. R., McCollom, T. M., Kiang, N. Y., and Jørgensen, B. B. (2023). The metabolic rate of the biosphere and its components. Proceedings of the National Academy of Sciences, 120(25):e2303764120.

[22] Hou, C., Zuo, W., Moses, M. E., Woodruff, W. H., Brown, J. H., and West, G. B. (2008). Energy uptake and allocation during ontogeny. Science (New York, N.Y.), 322(5902):736–739.

[23] Kleiber, M. (1932). Body size and metabolism. Hilgardia, 6(11):315–353.

[24] Kooijman, B. (2009). Dynamic Energy Budget Theory for Metabolic Organisation. Cambridge University Press, Cambridge, 3 edition.

[25] Koz-lowski, J., Konarzewski, M., and Gawelczyk, A. T. (2003). Cell size as a link between noncoding DNA and metabolic rate scaling. Proceedings of the National Academy of Sciences of the United States of America, 100(24):14080–14085.

[26] Labra, F. A., Bogdanovich, J. M., and Bozinovic, F. (2016). Nonlinear temperature effects on multifractal complexity of metabolic rate of mice. PeerJ, 4:e2607.

[27] Labra, F. A., Marquet, P. A., and Bozinovic, F. (2007). Scaling metabolic rate fluctuations. Proceedings of the National Academy of Sciences, 104(26):10900–10903.

[28] Legendre, L. J. and Davesne, D. (2020). The evolution of mechanisms involved in vertebrate endothermy. Philosophical Transactions of the Royal Society B: Biological Sciences, 375(1793):20190136.

[29] Maino, J. L., Kearney, M. R., Nisbet, R. M., and Kooijman, S. A. L. M. (2014). Reconciling theories for metabolic scaling. Journal of Animal Ecology, 83(1):20– 29.

[30] Mandelbrot, B. B. (1983). The fractal geometry of nature. New York.

[31] Marquet, P. A., Quiñones, R. A., Abades, S., Labra, F., Tognelli, M., Arim, M., and Rivadeneira, M. (2005). Scaling and power-laws in ecological systems. Journal of Experimental Biology, 208(9):1749–1769.

[32] McMahon, T. A. (1975). Using body size to understand the structural design of animals: quadrupedal locomotion. Journal of Applied Physiology, 39(4):619–627.

[33] Munoz, M. A. (2018). Colloquium: Criticality and dynamical scaling in living systems. Reviews of Modern Physics, 90(3):031001.

[34] Norin, T. and Metcalfe, N. B. (2019). Ecological and evolutionary consequences of metabolic rate plasticity in response to environmental change. Philosophical Transactions of the Royal Society B: Biological Sciences, 374(1768):20180180.

[35] Oliveira, A. C., Rebelo, A. R., and Homem, C. C. F. (2021). Integrating animal development: How hormones and metabolism regulate developmental transitions and brain formation. Developmental Biology, 475:256–264.

[36] Patterson, M. R. (1992). A mass transfer explanation of metabolic scaling relations in some aquatic invertebrates and algae. Science (New York, N.Y.), 255(5050):1421–1423.

[37] Rubner, M. (1883). Über den Einfluss der Körpergrösse auf Stoff-und Kraftwechsel. Pages: 536 Publication Title: Zeitschrift für Biologie Volume: 19.

[38] Seymour, R. S. (2001). Biophysics and physiology of temperature regulation in thermogenic flowers. Bioscience Reports, 21(2):223–236.

[39] Solé, R., Kempes, C. P., Corominas-Murtra, B., De Domenico, M., Kolchinsky, A., Lachmann, M., Libby, E., Saavedra, S., Smith, E., and Wolpert, D. (2024). Fundamental constraints to the logic of living systems. Interface Focus, 14(5):20240010.

[40] Solé, R. V., Manrubia, S. C., Benton, M., Kauffman, S., and Bak, P. (1999). Criticality and scaling in evolutionary ecology. Trends in ecology & evolution, 14(4):156– 160.

[41] Stanley, H. E., Amaral, L. A., Buldyrev, S. V., Gold-berger, A., Havlin, S., Leschhorn, H., Maass, P., Makse, H., Peng, C.-K., Salinger, M., and others (1996). Scaling and universality in animate and inanimate systems. Physica A: Statistical Mechanics and its Applications, 231(1–3):20–48.

[42] Swanson, D. L., Stager, M., Vézina, F., Liu, J.-S., McKechnie, A. E., and Amirkhiz, R. G. (2023). Evidence for a maintenance cost for birds maintaining highly flexible basal, but not summit, metabolic rates. Scientific Reports, 13(1):8968.

[43] Vicsek, T. (2001). Fluctuations and scaling in biology. Oxford University Press New York.

[44] West, G. B., Brown, J., and Enquist, B. (2000). Scaling in biology: patterns and processes, causes and consequences. Scaling in biology, 87:112.

[45] West, G. B. and Brown, J. H. (2004). Life’s universal scaling laws. Physics today, 57(9):36–42.

[46] West, G. B., Brown, J. H., and Enquist, B. J. (1997). A General Model for the Origin of Allometric Scaling Laws in Biology. Science, 276(5309):122–126.

[47] West, G. B., Brown, J. H., and Enquist, B. J. (2001). A general model for ontogenetic growth. Nature, 413(6856):628–631.

[48] West, G. B., Woodruff, W. H., and Brown, J. H. (2002). Allometric scaling of metabolic rate from molecules and mitochondria to cells and mammals. Proceedings of the National Academy of Sciences of the United States of America, 99(Suppl 1):2473–2478.

[49] White, C. R., Alton, L. A., Bywater, C. L., Lombardi, E. J., and Marshall, D. J. (2022). Metabolic scaling is the product of life-history optimization. Science, 377(6608):834–839.

[50] White, C. R., Phillips, N. F., and Seymour, R. S. (2005). The scaling and temperature dependence of vertebrate metabolism. Biology Letters.

[51] Yang, X., Heinemann, M., Howard, J., Huber, G., Iyer-Biswas, S., Le Treut, G., Lynch, M., Montooth, K. L., Needleman, D. J., Pigolotti, S., Rodenfels, J., Ronceray, P., Shankar, S., Tavassoly, I., Thutupalli, S., Titov, D. V., Wang, J., and Foster, P. J. (2021). Physical bioenergetics: Energy fluxes, budgets, and constraints in cells. Proceedings of the National Academy of Sciences of the United States of America, 118(26):e2026786118.

## References

[1] G. B. West, W. H. Woodruff, J. H. Brown, Allometric scaling of metabolic rate from molecules and mitochondria to cells and mammals. Proceedings of the National Academy of Sciences of the United States of America 99, 2473 (2002).

[2] A. M. Makarieva, et al., Mean mass-specific metabolic rates are strikingly similar across life’s major domains: Evidence for life’s metabolic optimum. Proceedings of the National Academy of Sciences 105, 16994 (2008).

[3] E. Ortega-Arzola, P. M. Higgins, C. S. Cockell, The minimum energy required to build a cell. Scientific Reports 14, 5267 (2024).

